# Quorum sensing signals from epibiont mediate the induction of bioactive peptides in mat-forming cyanobacteria *Nostoc*

**DOI:** 10.1101/2021.04.23.441229

**Authors:** Subhasish Saha, Paul-Adrian Bulzu, Petra Urajová, Jan Mareš, Grzegorz Konert, João Câmara Manoel, Markéta Macho, Daniela Ewe, Pavel Hrouzek, Jiří Masojídek, Rohit Ghai, Kumar Saurav

## Abstract

The regulation of oligopeptides production is essential in understanding their ecological role in complex microbial communities including harmful cyanobacterial blooms. The role of chemical communication between the cyanobacterium and the microbial community harboured as epibionts within its phycosphere is at an initial stage of research and little is understood about its specificity. Herein, we present insight into the role of a bacterial epibiont in regulating production of cyanobacterial oligopeptides microviridins, well-known elastase inhibitors with presumed anti-grazing effects, in an ecologically important cyanobacterial genus *Nostoc*. Heterologous expression and identification of specific signal molecules from the epibiont suggest the role of a quorum sensing-based interaction. Further, physiological experiments show an increase in microviridin production without affecting cyanobacterial growth and photosynthetic activity. Simultaneously, oligopeptides presenting a selective inhibition pattern provide support for their specific function in response to the presence of cohabitant epibionts. Thus, the chemical interaction revealed in our study provides an example of an interspecies signalling pathway monitoring the bacterial flora around the cyanobacterial filaments and induction of intrinsic species-specific metabolic responses.

**IMPORTANCE:** The regulation of cyanopeptide production beyond microcystin is essential to understand their ecological role in complex microbial communities, e.g. harmful cyanobacterial blooms. The role of chemical communication between the cyanobacterium and the epibionts within its phycosphere is at an initial stage of research and little is understood about its specificity. The frequency of cyanopeptide occurrence also demonstrates the need to understand the contribution of cyanobacterial peptides to the overall biological impact of cyanopeptides on aquatic organisms and vertebrates including humans. Our results shed light on the epibiont control of cyanopeptide production via quorum sensing mechanisms and we posit that such mechanisms may be widespread in natural cyanobacterial bloom community regulation.

## OBSERVATION

Despite the rise in cyanobacterial bloom occurrence and detection of cyanopeptides (CNPs) beyond microcystin (1, 2) across the world, why and how of the regulation of these metabolites remains poorly understood (3). It has mostly been argued that the peptide net production rates are linearly correlated with the growth rate of the cyanobacterial cells, while a direct impact of environmental factors on peptide production is of relatively minor importance (4). However, the role of bacterial-cyanobacterial interactions on physiological control of CNPs production has never been evaluated. The majority of CNP producers frequently form biofilms and their associated epibionts might underlie secondary metabolite production as well as biofilm development. Microbial communities associated with cyanobacterial biofilms are attracted and held together by cohesive exopolysaccharide envelopes that can harbour numerous co-existing microbial species belonging to diverse lineages (5, 6). A recent study provided new insights into the role of quorum sensing (QS) signal molecules in cyanobacterial aggregation processes and biofilm development (7). Genes responsible for their synthesis/regulation could not be confidently identified in their genomes, which led us to speculate that cyanobacteria may have evolved a different mechanism for regulation of QS autoinducers and might rely on epibionts to produce them.

Single-filament picking of strain *Nostoc* sp. TH1SO1 (strain TH1SO1) followed by *de novo* genome sequencing and metagenomic binning allowed the recovery of one high-quality *Nostoc* metagenome-assembled genome (MAG), together with a total of five medium-to-high quality epibiont bins assigned to phyla Proteobacteria (n=3) and Bacteroidota (n=2) (Supplementary Table S1), as well as three low-quality bins derived from Proteobacteria (bins 5, 6 and 9; https://figshare.com/s/817256304aa3f038bd85) (Supplementary methods). The draft genome of strain TH1SO1 was retrieved in 247 genomic contigs amounting to 7,653,454 bp (99.56% estimated completeness, 0.3% contamination) (Supplementary Table S1). Three complete putative biosynthetic gene clusters (BGCs) for microviridins (MDNs), a ribosomally synthesised and post-translationally modified peptide (RiPP) containing five unique functional precursor peptides (MdnA) were found in the strain TH1SO1 genome (Fig. 1A-B). One of these gene clusters (cluster-a), for which we have detected the predicted products, encoded three functional MDN precursor genes, both lactam and lactone macrocycle-forming ATP-grasp ligases, an *N*-acetyltransferase, and an ABC-type transporter (Fig. 1A).

**Fig. 1.**
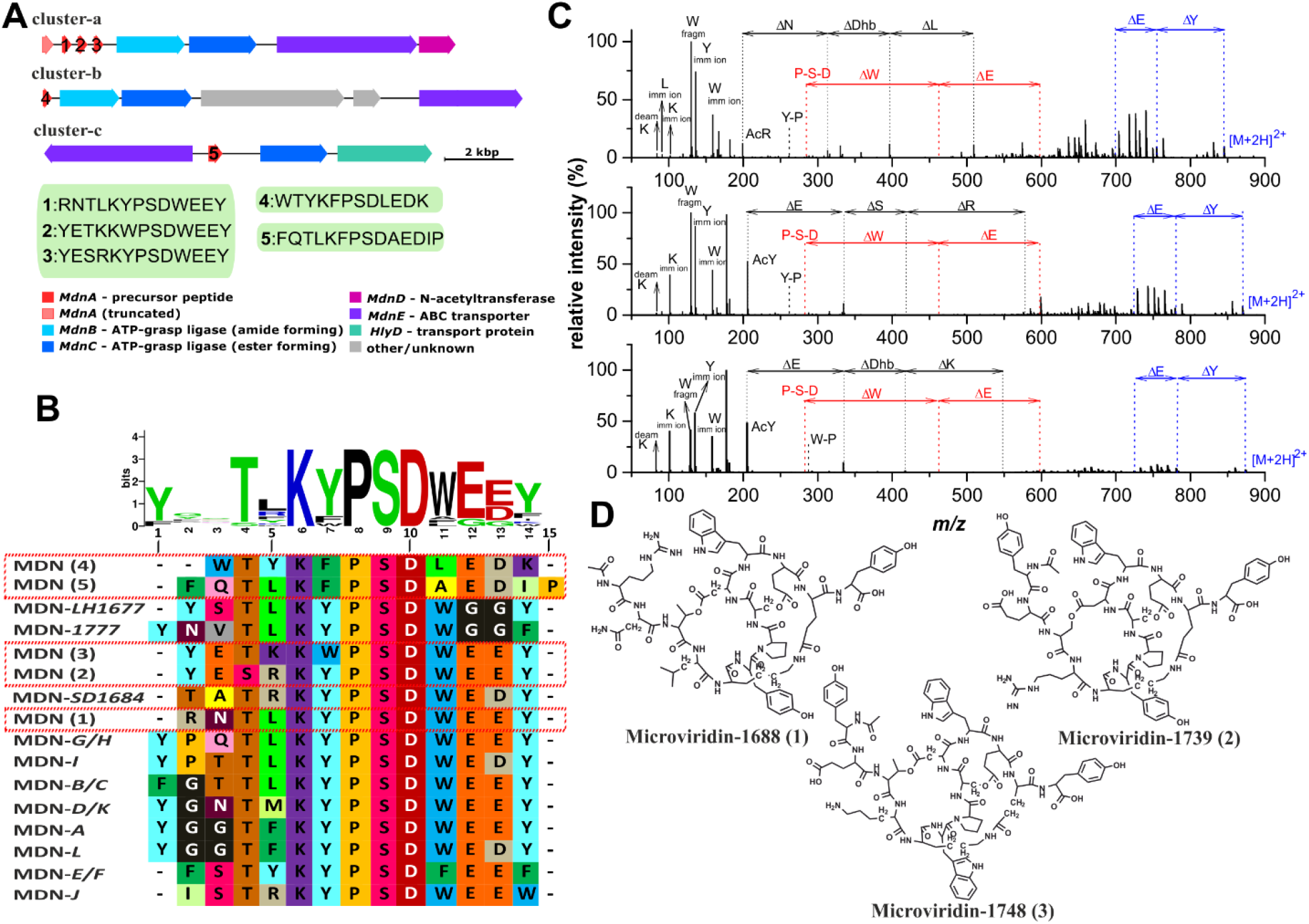
**A**. Gene map of microviridin gene clusters (a-c) mined from the *Nostoc* sp. TH1SO1 genome. a, three functional precursor peptides (MdnA) differing in the core peptide sequence are predicted, whereas for cluster b and c, each encode a single precursor peptide – core peptide sequences are indicated in green boxes. **B**. Variation in microviridin peptide sequence, multiple sequence alignment detected all the five microviridin precursors as novel and differing in from 2 to 6 amino acid positions compared to the known variants. The consensus sequence revealed the variation in conserved motif KYPSD where Y (Tyr) was replaced by F (Phe) and W (Trp). **C**. HR-MS/MS product ion spectra of protonated microviridins from *Nostoc* sp. TH1SO1, **D**. Structure of the three detected microviridins; Microviridin-1688 (*m*/*z* 844.8917 [M + 2H]^+^), Microviridin-1739 (*m/z* 870.3707 [M + 2H]^+^) and Microviridin-1748 (*m/z* 874.8821 [M + 2H]^+^) confirmed by coupling product ion spectra and the genomic data.

Fractionation of the biomass extracts and manual inspection of the spectra acquired using HPLC-HRMS/MS suggested the presence of three novel MDNs originating from gene cluster-a. Subsequent chromatographic separation led us to isolate Microviridin-1688 (*m*/*z* 844.8917 [M + 2H]^2+^ (1.2 mg)), Microviridin-1739 (*m/z* 870.3707 [M + 2H]^2+^ (0.3 mg)) and Microviridin-1748 (*m/z* 874.8821 [M + 2H]^2+^ (1.1 mg)) in pure state and their structures were predicted and corroborated by comparing the genomic data (sequence of the encoded core peptide) to the product ion mass spectra (Fig. 1A-D; Supplementary Table S2). The predicted products from gene clusters-b and c were not detected, a possible explanation for this being these BGCs were silent in current culture conditions.

It has been postulated that factors such as buoyancy regulation or interaction with grazers or pathogens promote the differentiation among cyanobacterial chemotypes (8). This prompted us to investigate the BGCs of the recovered epibiont bins. Four autoinducer synthase gene clusters were detected and among them one of the autoinducer synthase, *SGBI* (630 bp) belonging to the genus *Sphingobium (contig-2686)* was successfully heterologously expressed in BL21(DE3)/pET28a (Fig. 2A). Monitoring of the characteristic product ion at *m/z* 102.0550 (9) confirmed the presence of six homoserine lactones (HSLs) possessing hydroxylated fatty acyl side chains of different length (3-hydroxy-C_7_-HSL, 3-hydroxy-C_8_-HSL, 3-hydroxy-C_9_-HSL, 3-hydroxy-C_10_-HSL, 3-hydroxy-C_12_-HSL, and 3-hydroxy-C_14_-HSL) (Fig. 2B-C; Supplementary Fig. S1; supplementary methods). The presence of HSLs during cyanobacterial blooms has been reported, with their concentrations reaching up to 10 mg/L (10-12). The role of epibionts on microalgal growth and QS-dependent physiological regulations has already been established using transcriptomic approach in marine environment (13). To assess the role of QS-dependent regulation on CNPs production, we mimicked the bacterial load in the culture of strain TH1SO1 with two major variants of HSLs (3-hydroxy-C_8_-HSL and 3-hydroxy-C_10_-HSL) at 2.5 µM final concentration (Supplementary methods). A significant increase of up to two-fold in the production of the MDN-1688 (once normalised to dry biomass) (Fig. 2D) without any difference in photosynthetic activity (supplementary Fig. S2; supplementary methods) among the control and fed-batch cultures was observed. In a similar experimental set up, MDNs production was reduced or unchanged when fed with 3-hydroxy-C_8_-HSL/3-hydroxy-C_10_-HSL together with a known QS inhibitory molecule, penicillic acid (Fig. 2E), suggesting that these results are not artefacts of an unexplained mechanism like inhibition or chemotype specificity. These results encouraged us to study the role of MDNs in maintaining the niche colonisation or facilitating the biofilm formation within the phycosphere. For example, MDN-J isolated before was shown to inhibit the molting process of *Daphnia*, providing an advantage in maintaining and survival of the dense community during bloom formation (14). Similarly, for sustained colonisation by specific taxa, an intricate mechanism must be involved with localised specificity. Our results showed that MDNs inhibited (32-55%) the bioluminescence of QS bioreporter strain *E. coli* pSB1075 (*LasR*-based) in a dose-dependent manner (Fig. 2F). In contrast, no inhibition of bioluminescence on *E. coli* pSB401 (*LuxR*-based) was observed for MDNs. The inhibition of bioluminescence against one of the reporter strains at non-inhibitory concentration suggests its specificity in inhibiting *lasR* system implying its possible role in monitoring the selection of a specific epibiont colonising using *luxR*-based QS system.

**Fig. 2.**
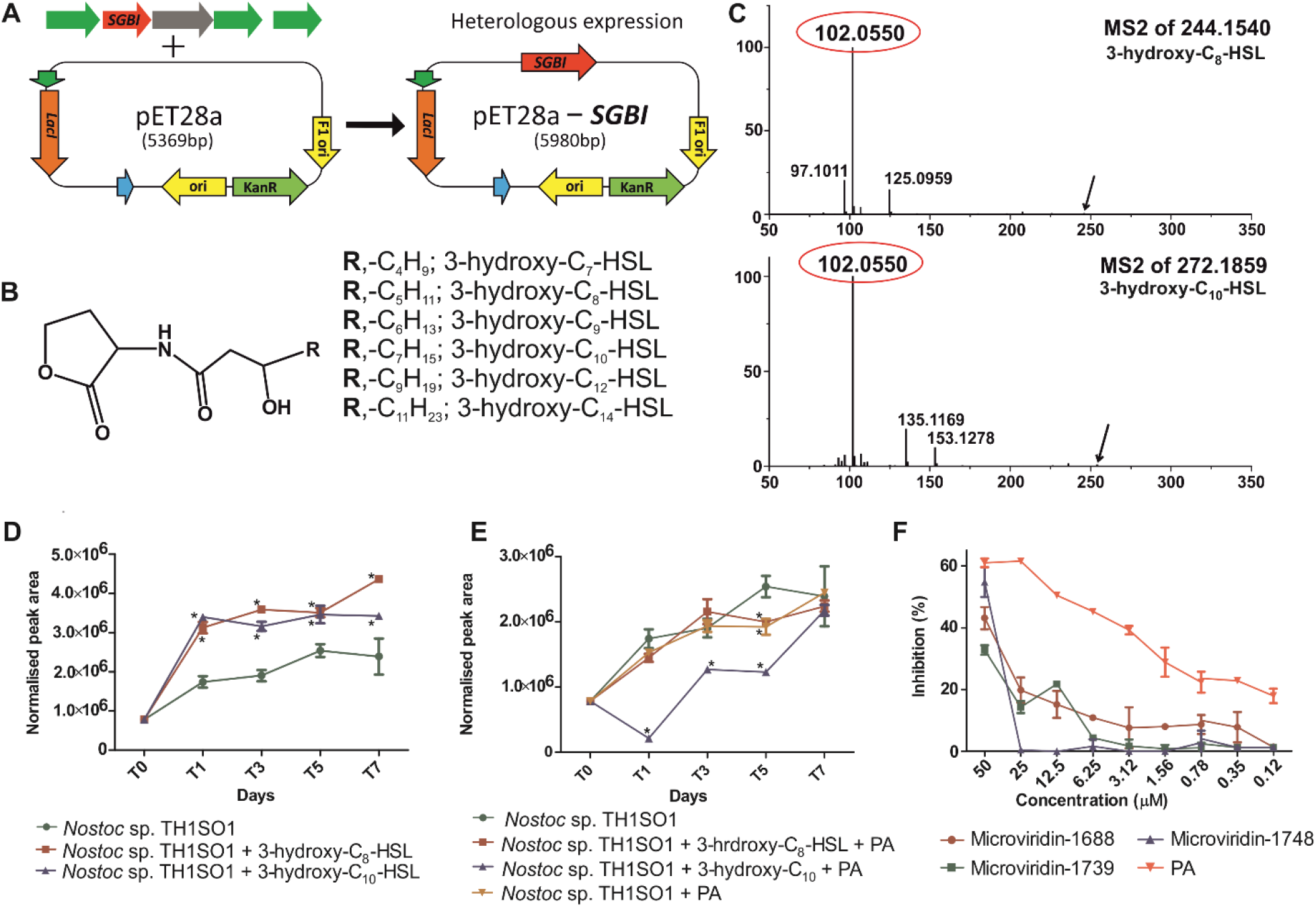
**A**. Heterologous expression of *SGBI* in BL21(DE3)/pET28a; **B**. General structure of all the variants of HSL detected from the extract of heterologously expressed *SGBI* in BL21(DE3)/pET28a; **C**. HR-MS/MS product ion spectra of two most abundant protonated molecule [M + H]^+^) at *m/z* 244.1540 (3-hydroxy-C_8_-HSL) and *m/z* 272.1859 (3-hydroxy-C_10_-HSL) derived from the extract of heterologously expressed *SGBI* in BL21(DE3)/pET28a. The characteristic product ion at *m/z* 102.0550, corresponding to the deacylated homoserine lactone, was detected; **D**. Induction of Microviridin-1688 production after feeding with 3-hydroxy-C_8_-HSL/3-hydroxy-C_10_-HSL (at 2.5 µM final concentration), the LC-MS peak area was normalized to dry biomass; **E**. Inhibition of Microviridin-1688 production in the presence of a quorum sensing inhibitor, penicillic acid (PA) in combination with 3-hydroxy-C_8_-HSL/3-hydroxy-C_10_-HSL (at 2.5 μM final concentration) and **F**. Dose-dependent inhibition activity of Microviridin-1688, Microviridin-1739, Microviridin-1748 and penicillic acid on QS-dependent bioluminescence of the *LasR*-based bioreporter strain *E. coli* pSB1075 induced by its cognate molecule 3-oxo-C_10_-HSL at non-inhibitory concentration. The average bioluminescence observed for the negative control is used to calculate relative inhibition percentage. Data are expressed as SD of mean (*n* = 3). * *P* < 0.001 versus control by ANOVA followed by Bonferroni post-test.

Our results have demonstrated the potential role of QS in the regulation of CNPs production and show that these processes might play a key role in epibiont-cyanobacterial interactions. It further highlights the value of culture-based experimentation and the importance of developing a model organism for studying complex ecological interactions. This knowledge could in turn permit a bottom-up reconstruction of the multipartite interactions, mediated by the exchange of secondary metabolites (15, 16). Key to this exchange are the regulatory circuits that control the induction of secondary metabolites (16). The question of when and how a bacterium ‘chooses’ to induce a given BGCs is a fascinating and unresolved one, and detailed studies are sure to illuminate mechanisms, perhaps new ones, by which exogenous signals govern this process.

## Data availability

Sequence data generated during this study have been deposited in NCBI under and linked to BioProject ID PRJNA718890. Genome bins assembled in this study have been deposited at DDBJ/ENA/GenBank under the accessions JAGKSW000000000-JAGKTB000000000. The version described in this paper is version JAGKSW010000000. Individual cotings encoding genes of interest discussed within this study are available under GenBank accessions MW857567-MW857570. The derived data that support the findings of this paper, including the assembled metagenomic contigs collection, are available in figshare with the identifier https://figshare.com/s/817256304aa3f038bd85.

